# A p21-GFP zebrafish model of senescence for rapid testing of senolytics *in vivo*

**DOI:** 10.1101/2022.09.19.506911

**Authors:** Samir Morsli, Catarina M. Henriques, Pamela S Ellis, Heather Mortiboys, Sarah Baxendale, Catherine Loynes, Stephen A. Renshaw, Ilaria Bellantuono

## Abstract

Senescence drives the onset and severity of multiple ageing-associated diseases as well as frailty. As a result, there has been an increased interest in mechanistic studies and in the search for compounds targeting senescent cells, known as senolytics. Mammalian models are commonly used to test senolytics and generate functional and toxicity data at the level of organs and systems, yet this is expensive and time consuming. Zebrafish share high homology in genes associated with human ageing and disease. They can be genetically-modified relatively easily. In larvae, most organs develop within 5 days of fertilisation and are transparent, which allows tracking of fluorescent cells *in vivo* in real time, testing drug off-target toxicity and assessment of cellular and phenotypic changes. Here, we have generated a transgenic zebrafish line that expresses green fluorescent protein (GFP) under the promoter of a key senescence marker, p21. We show an increase in p21:GFP^+^ cells in larvae following exposure to ionising radiation and with natural ageing. p21:GFP^+^ cells display other markers of senescence, including senescence-associated β-galactosidase and IL6. The observed increase in senescent cells following irradiation is associated with a reduction in the thickness of muscle fibres and mobility, two important ageing phenotypes. We also show that quercetin and dasatinib, two senolytics currently in clinical trials, reduce the number of p21:GFP^+^ cells, in a rapid 5-day assay. This model provides an important tool to study senescence in a living organism, allowing the rapid selection of senolytics before moving to more expensive and time-consuming mammalian systems.

## Introduction

Senescent cells are characterised by cell-cycle arrest, loss of function (despite persisting metabolic activity) and by the secretion of multiple pro-inflammatory and tissue-remodelling factors, known as the the senescence induced secretory phenotype (SASP). Senescence can be triggered by internal stimuli, including persistent DNA damage, telomere dysfunction or oncogene activation or external stimuli, such as ionising radiation [1]. In animal models, the burden of senescent cells increases with age in multiple tissues [2, 3], while their elimination improves tissue homeostasis with age, preventing the onset or limiting the severity of multiple age-associated diseases [4]. Senescent cell clearance, therefore, offers great promise for the prevention of multimorbidity and frailty, two of the biggest challenges for modern healthcare and as such, has galvanised interest in developing new drugs to reduce the burden of senescent cells.

However, there are major challenges impeding mechanistic studies and testing of compounds to reduce senescent cell burden. *In vitro* systems do not provide the same level of information on toxicity, cell-cell and organ-organ interaction as animal models, yet the use of *in vivo* mammalian systems takes significant time and resources. Zebrafish models can often bridge the gap between *in vitro* systems and *in vivo* mammalian systems. Zebrafish share 84% of known human disease-associated genes and 70% of human protein encoding genes [5] and have unique features as a model organism. They are born in large clutches and grow quickly, rapidly providing high numbers at low cost. Larval zebrafish are almost completely optically transparent and amenable to genetic manipulation, allowing ready generation of fluorescent transgenic reporters to track individual cells within the same live animal, over time. By combining different fluorescent reporters, it becomes possible to image the interaction between cell types and elucidate novel mechanisms, as has been shown with labelling of cells of the innate immune system [6, 7]. Numerous zebrafish models of disease are available and several have been shown to be amenable to studying drug efficacy *in vivo* [8]. Zebrafish larvae fit in 96-well plates and can be assessed for potential off-target toxicity and cellular and phenotypic changes across multiple tissues, using assays which last only a few days. For example, with automated quantitation, more than 500,000 zebrafish larvae were screened to identify novel compounds that increase the number of insulin-producing β-cells in the pancreas, as a potential treatment for diabetes [9].

In this study we report the generation of a transgenic zebrafish line *TgBAC(cdkn1a:GFP)sh506* (termed p21:GFP thereafter), which expresses Green Fluorescent Protein (GFP) under the promoter of a key senescence marker, p21 (*cdkn1a*). p21 is one of the major regulators and cellular markers of senescence [10]. Recent reports show that cells expressing high levels of p21 *in vivo* express other markers of senescence and their numbers increase with age, in multiple murine organs [11]. In addition, elimination of p21^+^ senescent cells reduces frailty and attenuates insulin resistance in obese mice [12], while elimination of p21^+^ cells, but not p16^+^ cells, improves radiation-induced osteoporosis [13], suggesting that p21 is a good marker of senescent cells. Zebrafish have a homologue of the human p21 gene, known to increase with increased levels of p16-like expression and senescence associated beta-galactosidase [14].

Here, we demonstrate that expression of p21-GFP is upregulated in larvae following exposure to ionising radiation and with natural ageing. Importantly, accumulation of senescent cells is associated with reduction in muscle fibre thickness and mobility, two important ageing phenotypes. We have identified a population of cells that express high levels of p21:GFP (p21:GFP^bright^) and co-express other markers of senescence, including IL6, a major SASP factor. Finally, we show that the p21:GFP transgenic zebrafish line can be used for imaging and serves as a useful readout of senescence when testing for the most effective senolytic drugs *in vivo*, using 96-well plates in a rapid 5-day assay. This line, therefore, provides an important tool to study senescent cells in a living organism, allowing for the rapid testing of drugs, before moving to more expensive and time-consuming mammalian systems. Importantly, the use of zebrafish less than 5 day post-fertilization (dpf) fulfil the principle of the 3Rs as experiments at this early stage are not categorised as animal studies by regulatory bodies and their use could replace high-order animals.

## Results

### Zebrafish larvae show markers of senescence following ionising radiation

To determine whether it was possible to induce senescence in zebrafish larvae, we exposed the larvae to 12 Gy irradiation at 2 dpf (Fig 1a). These parameters were chosen because they were the highest dose and the earliest time point possible that did not result in an abnormal development or reduced viability (extended data Fig 1). Abnormal development was assessed by counting the number of abnormalities (scored as pericardial oedema, deflated swim bladder, spinal curvature, skin lesions and stunted growth) in 140 fish over 3 experiments. The percentage of abnormalities scored in the non-irradiated group was 10.2±6.6%, with no significant difference across the 3 days. There were significantly more abnormalities by 5dpf in zebrafish larvae irradiated at 1dpf (64±13.2%). However, the number of abnormalities by 5dpf were in a similar range to non-irradiated zebrafish larvae when the irradiation was performed at day 2 (15±4.6%) or at day 3 (19.6±8.7). Therefore, we chose to irradiate at 2dpf from here on. Multiple markers of senescence were assessed at 5dpf (Fig 1a). We observed a significant increase in the expression of mRNA for *p21* (p=0.0005, n=4), *p16-like* (*cdkn2a/b)(* p=0.0002, n=4), *p53* (P=0.0002, n=4), *cyclinG1* (p=0.008, n=3), and SASP factor *MMP2* (p=0.0005, n=4) by qPCR, using whole fish RNA extracts from 50 pooled larvae in each experiment (Fig 1b). No significant increase in the levels of SASP marker *IL8* was observed (Fig 1b). To determine the spatial location of senescent cells, we performed whole-mount *in situ* hybridisation using a previously published p21 probe [15]. We observed a significant increase in p21 expression following irradiation, which increased in a dose-dependent manner (Fig 1c). This was particularly strong in the pharyngeal arches, brain and intestinal regions of the zebrafish and overlapped with areas where there was an increase in senescence-associated βGal expression (SA-βGal) (Fig 1d). Immunostaining for γH2AX DNA damage foci, another marker of senescence, showed a significant increase in the percentage of cells with bright nuclear γH2AX staining (individual foci were not distinguishable) in the ventral head region, following irradiation (Fig.1e). These ranged from 3 to 10%, comparable to the levels seen in aged organisms [16–18].

**Figure 1.**
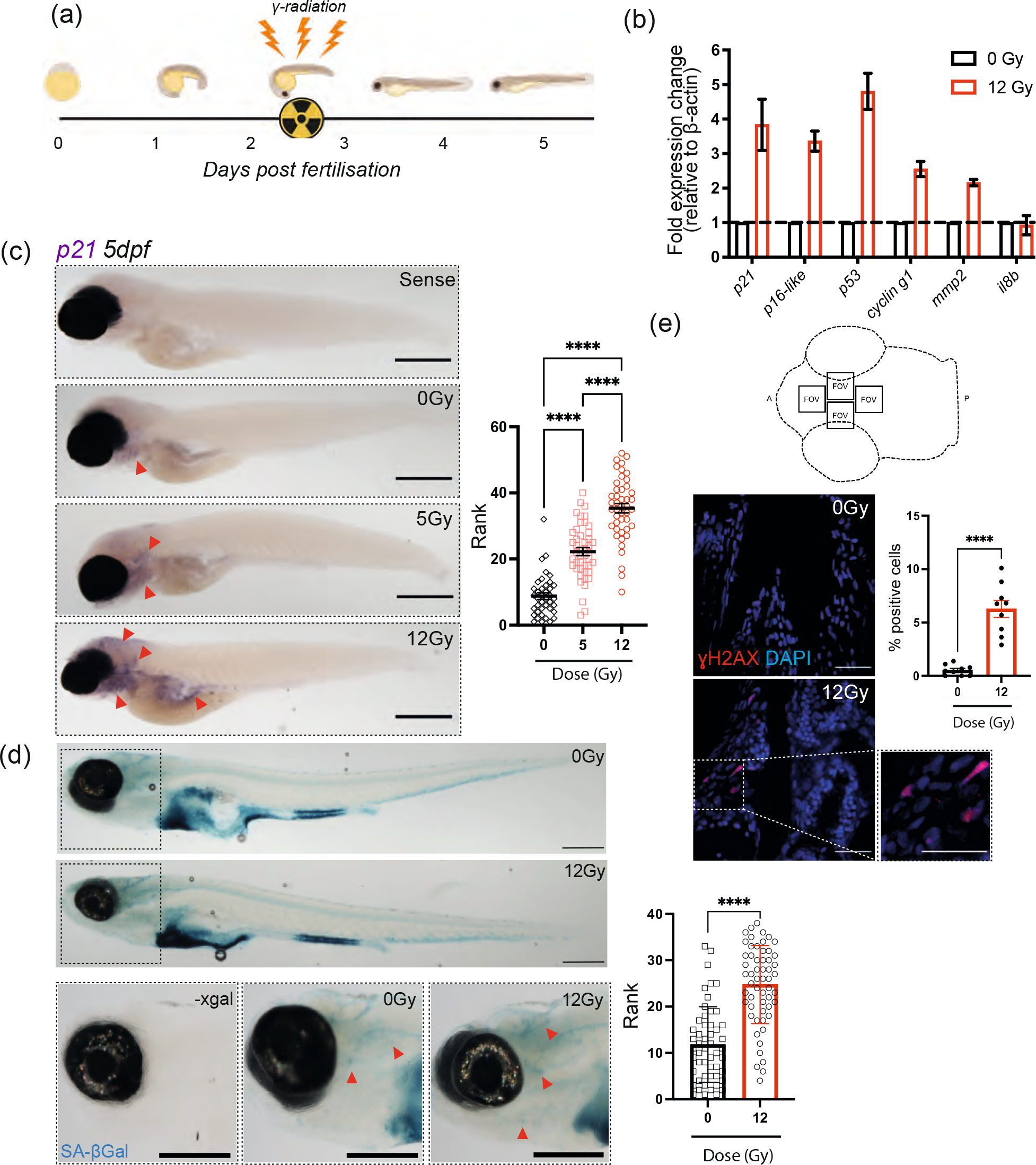
Irradiation of zebrafish larvae upregulates multiple markers of senescence. (a) Diagram depicting the experimental protocol used to induce senescence in zebrafish larvae using Cs_137_ ɣ-Irradiation at 2dpf and assessing markers of senescence at 5dpf. (b) Quantitative PCR (qPCR) of whole zebrafish mRNA at 5dpf following 12Gy irradiation to determine gene expression of *p21 (cdkn1a), p16-like (cdkn2a/b), p53, cyclin-g1, mmp2, IL8b*. Fold expression was calculated by 2^-ΔCt relative to β-actin. mRNA was pooled from 50 zebrafish for each independent repeat. The graph represents the mean ± SEM of 3 repeats. (c) Transmitted light photomicrographs of a whole-mount *in-situ* hybridisation (WISH) for *p21* (*cdkn1a)* mRNA expression at 5dpf following 5 or 12Gy irradiation at 2dpf (left panels). Areas of increased staining include pharyngeal arches, brain, and intestine, depicted with red arrows. Scale 250μm. Quantification of ISH photomicrographs through blind ranking such that fish with the strongest staining are ranked highest (right panel). Data were examined by Kruskal Wallace test with Dunn’s multiple comparisons (N=45). (d) Photomicrographs of zebrafish at 5 dpf stained for senescence-associated β-Galactosidase (SA-βGal) activity. Representative examples of the head region are also displayed on the left. Scale 200μm (Left panels). Quantification of SA-βGal activity in the head region by blind ranking is on the left (n=55)(right panel). Data were examined by Kruskal Wallace test with Dunn’s multiple comparisons (e) Confocal fluorescence photomicrographs (Scale 25μm) and quantification of ɣH2AX immunofluorescence in the ventral zebrafish head regions (areas depicted in the cartoon in the top panel) a minimum of 600 cells/fish were counted. For each fish, 4 fields of view and 3 individual z planes, 10μM apart were analysed. At least 9 fish were analysed across three independent repeats. Data represented as mean ± SEM (N=9) and examined by unpaired t-test. Significant differences displayed as ** *p* < 0.01; **** *p* < 0.0001. a, Anterior; p, posterior; FOV, field of view.

### Irradiated larvae show signs of muscle wasting similar to aged zebrafish

One of the characteristics of ageing across organisms, including humans, and one of the main signs of frailty, is loss of body mass associated with muscle wasting [19]. In order to assess whether irradiation caused changes in muscle phenotype, we performed histological analysis of muscle in the ventral region of zebrafish larvae at 12 dpf following 12Gy of irradiation. There was a significant decrease in muscle fibre thickness compared to the non-irradiated control. This was similar to the decrease observed with natural ageing in the muscle of middle-aged (18 months) and geriatric fish (>36 months) (Fig 2a). More importantly, when larvae were placed individually in a 24-well plate and their movement quantified over a 30-minute period, at 5 and 12 dpf, we observed a significant reduction in the distance travelled at both time-points in irradiated larvae, compared to non-irradiated larvae (Fig. 2b). These data suggest that irradiated larvae develop similar muscle changes to aged zebrafish and that histological changes in muscle fibre thickness were accompanied by loss of muscle function.

**Figure 2.**
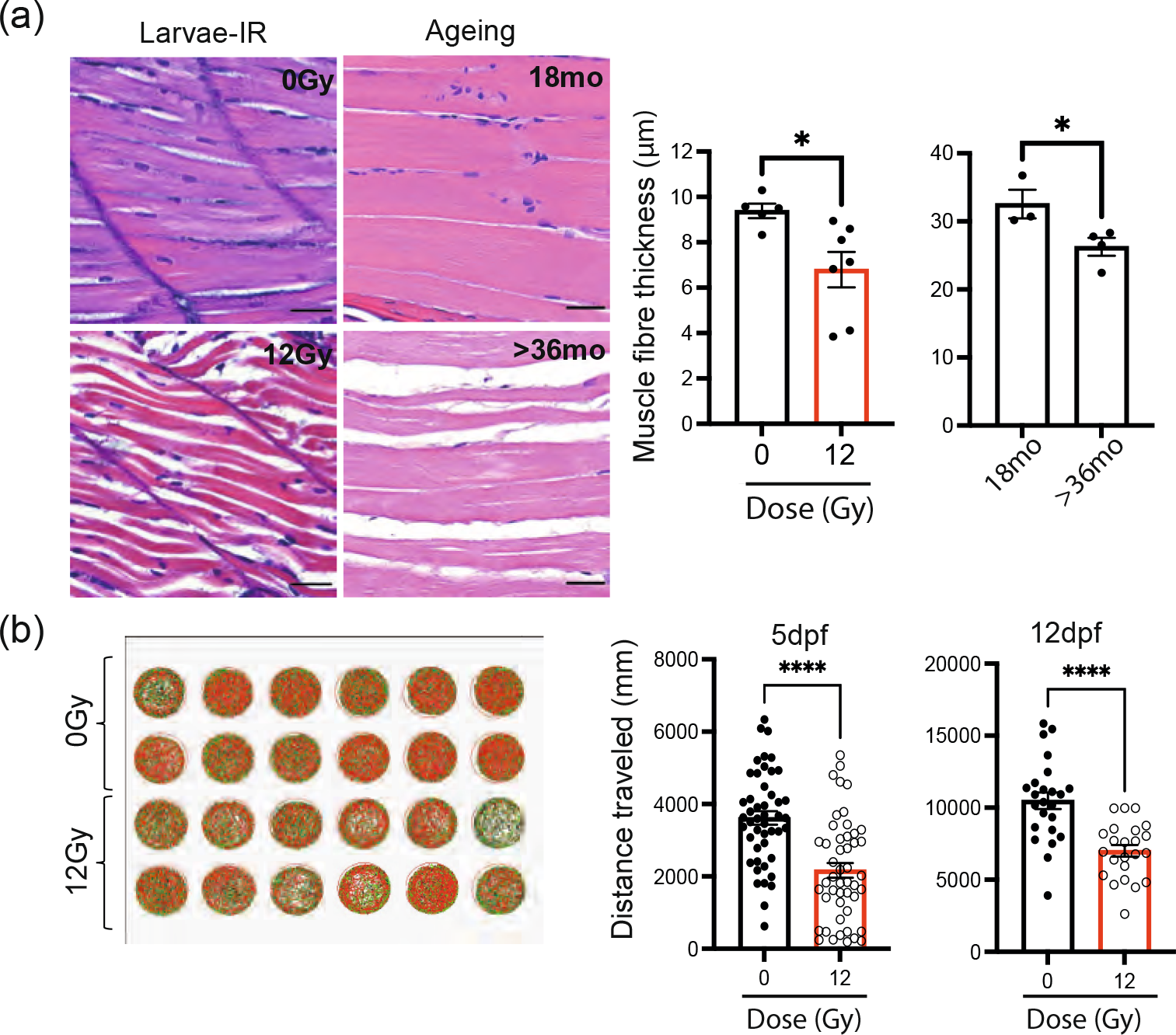
Zebrafish larvae showed reduced muscle fibre thickness and mobility following irradiation. (a) Representative photomicrographs of haematoxylin & eosin (H&E) staining of zebrafish muscle from either 12dpf larvae with and without irradiation (top left panel), and middle aged and geriatric adults (bottom left panel) (scale 25μm) and quantification of muscle fibre thickness in larvae following irradiation and in middle aged (18 months) and geriatric zebrafish (>36 months)(right panels). Data shown as mean ± SEM. Each dot represent an animal. (b) Representative example of distance travelled by zebrafish over 30 minutes in a 24-well plate at 5 (N ≥ 47) and 12 dpf (N ≥ 23) following 0Gy or 12Gy irradiation administered at 2dpf and quantitation of the distance travelled at 5 and 12 dpf. Each dot represents an animal. Data examined by unpaired t test. **** *p <0.0001;* * *p* < 0.05.

### Generation of a p21:GFP zebrafish model

To generate a p21:GFP reporter transgenic line, we used the DKEY 192-O24 bacterial artificial chromosome (BAC) encompassing the *p21* locus and containing at least 100kbp downstream and 50 kbp upstream of the start codon, to include as much of the promoter of the gene as possible. The BAC was modified such that the GFP sequence was incorporated into the BAC at the initial ATG start codon of the *p21*-encoding region in the first exon, common to both *p21* splice variants, and thereby placed under regulation of the *p21*-promoter (Fig 3a). The insertion of GFP disrupted the expression of the *p21* gene contained in the BAC and ensured it did not express an extra copy of *p21* when inserted into the zebrafish genome. To improve the likelihood that the modified BAC was incorporated into the zebrafish genome, a transposon-mediated system was used [20] (Fig 3b-c).

**Figure 3.**
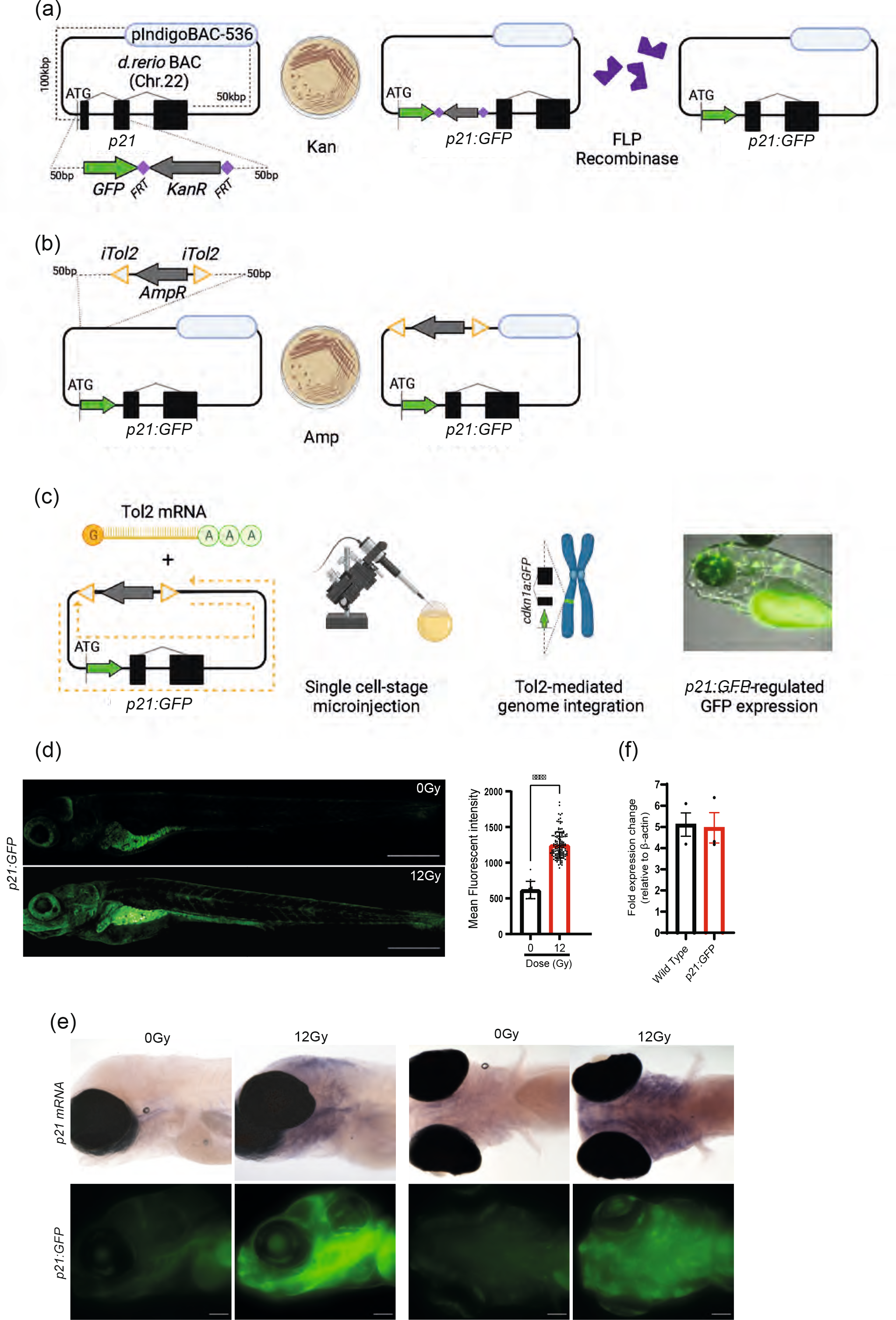
Generation of p21:GFP transgenic fish. (a-c) Diagram depicting development of TgBAC(*p21*:*GFP*)sh506 transgenic zebrafish (p21:GFP). (a) Identification of the DKEY 192-O24 Bacterial Artificial Chromosome (BAC) containing the *p21 l*ocus and modification with GFP cassette after the initial start codon of *p21*. Positive selection was carried out by identifying bacterial colonies expressing the kanamycin resistance cassette, which was later removed from the BAC with FLP FRT recombination. (b) Modification of the GFP-BAC with tol2-recognition sequences allowed tol2-mediated transgenesis. (c) Injection of *p21:GFP*-tol2-BAC alongside tol2 mRNA into fertilised single-cell-stage zebrafish embryos to create transgenic zebrafish reporter for *p21* promoter regulation. Founder zebrafish were identified according to *p21* promoter-associated GFP expression in their offspring, confirming germline expression of the transgene. (d) Representative confocal fluorescence photomicrographs to depict GFP fluorescence of 5dpf *p21:GFP* zebrafish following 0Gy or 12Gy irradiation at 2dpf (Scale 500μm) and quantification of fluorescence intensity of the whole p21:GFP transgenic zebrafish. Each dot represent an animal. Graph represents Mean±SEM and data were examined by Mann-Whitney test. (e) Transmitted and wide-field fluorescence photomicrographs taken laterally and ventrally showing p21:GFP fluorescence recapitulated endogenous *p21* mRNA expression (Scale bar 100μm). (f) qPCR of whole zebrafish mRNA demonstrating *p21* expression in transgenic *p21:GFP* and wild-type strain zebrafish, relative to βactin (2^-ΔCt). mRNA was pooled from 50 zebrafish for each independent repeat. The graph represent the mean ± SEM of 3 repeats. Data were examined by 2way ANOVA with Tukey’s multiple comparison test. **** *p < 0.0001;* *** *p* < 0.001; ** *p* < 0.01.

To verify that GFP expression was regulated in a similar way to endogenous p21, we measured green fluorescence intensity following 12Gy irradiation. An increase in mean fluorescent intensity was observed post-irradiation, which was more pronounced in the intestine, head regions and pharyngeal arches (Fig 3d), similar to that observed using *in situ* hybridisation and a p21 probe (Fig 3e). To verify that the transgenic reporter had comparable endogenous *p21* expression to wild-type fish and there was no additional contribution from the BAC, assessment of *p21* mRNA expression levels was performed by qPCR in the whole p21:GFP zebrafish at 3 days post-irradiation. As expected, we found an increase in p21 expression with irradiation in both wild-type and transgenic animals but there was no difference in levels of expression between the two lines (Fig 3f), suggesting that there was no additional p21 expression due to the presence of the BAC.

### The number of p21:GFP^bright^ cells increases with irradiation and natural ageing

To quantify the number of p21:GFP^+^ cells, flow cytometric analysis of zebrafish larvae post-irradiation was performed at 5 and 12dpf using the gating strategy shown in Fig 4a-c. Before irradiation we noticed a population of p21:GFP^dim^ cells at 5dpf (23.3%±3.4%, n=3)(Fig 4b), which did not significantly increase following irradiation when compared to the non-irradiated zebrafish larvae and remained consistent at later time points. In contrast, we detected the appearance of a GFP^bright^ population of cells at 5dpf following irradiation (0.95%±0.01% vs 14.06%±1.01% 0 and 12Gy respectively, n=3 p<0.05)(Fig.4b). The presence of this GFP^bright^ population persisted in the irradiated zebrafish larvae at 12 dpf (1.03%±0.05% vs 6.65%±0.72% 0 vs 12Gy respectively, n=3, p<0.05) although at lower levels than that observed at 5dpf (Fig 4c). Recently, Wang et al. (2022)[10] also reported a population of p21GFP^high^ cells accumulating in multiple tissues with age using an inducible p21-cre GFP mouse model. Indeed, analysis of brain, intestine and liver of middle-aged and geriatric zebrafish also showed a significant increase in p21:GFP^bright^ cells whereas no statistical difference was observed in the p21:GFP^dim^ cells (Fig 4c). Notably, tissues showed time and tissue-specific increases in the accumulation of senescent cells with age. Whilst the intestine showed a significant increase at middle age, the brain showed a more progressive increase through the ages and the liver seems to suffer a significant increase mainly at geriatric age (Fig 4c). This is in line with what was reported by Henriques *et al* (2013) [21]and Carneiro *et al*. (2016) [22] where a time and tissue-specific degeneration was observed with ageing wild type zebrafish and accelerated in the prematurely aged telomerase mutant (*tert^−/-^*) zebrafish, with the intestine as one of the first tissues to degenerate and accumulate senescence.

**Figure 4.**
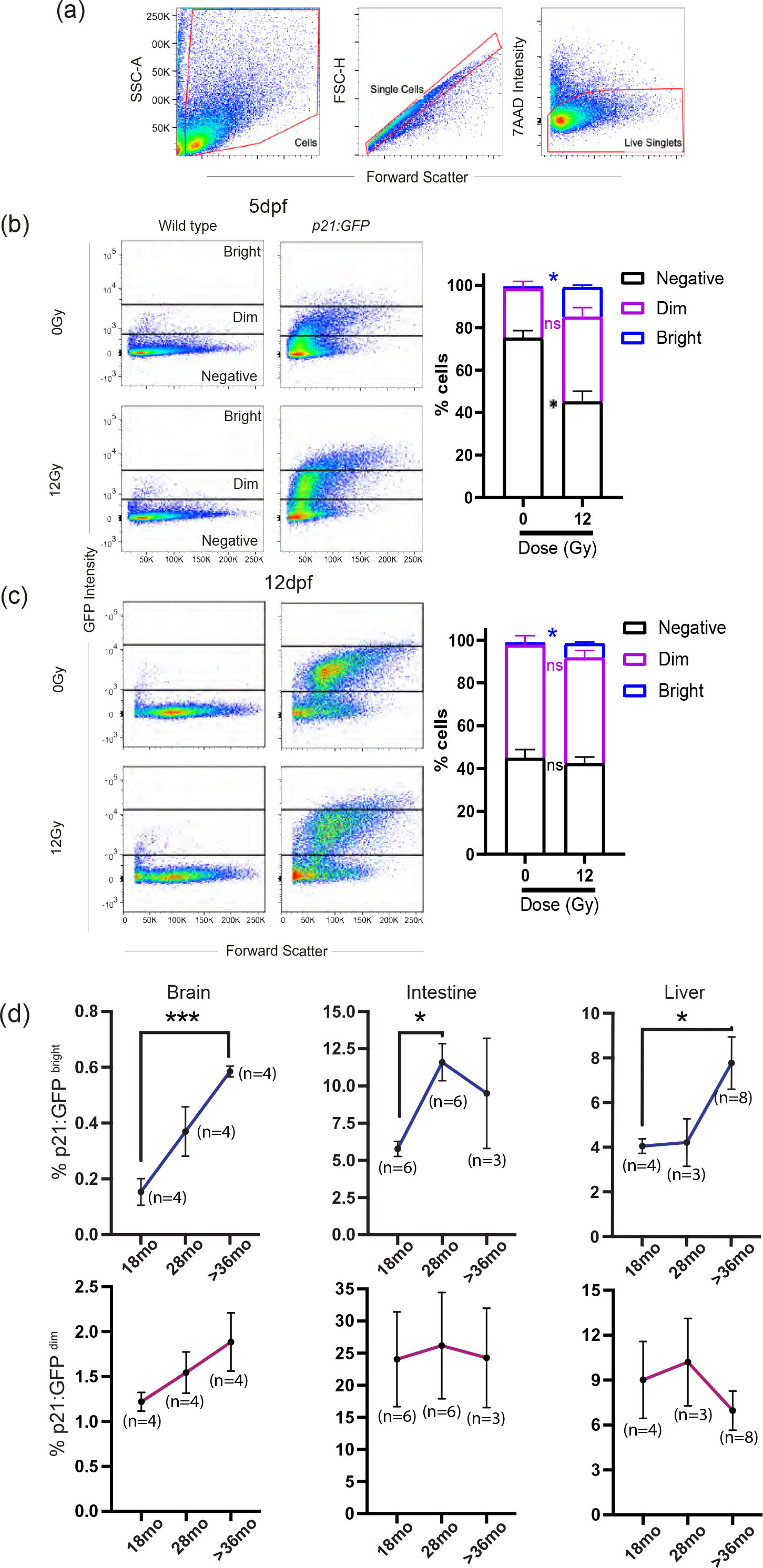
*p21:GFP*^Bright^ cells are induced with irradiation and ageing. (a) A representative example of the gating strategy used to assess GFP intensity by flow cytometry. Dissociated cells were gated to remove debris using forward and side scatter (left panel). The population was then gated for single cells using FS-H, FS-A (middle panel). Live cells were gated as 7-Aminoactinomycin D (7AAD) negative cells (right panel). Live cells were then assessed for p21:GFP intensity as shown in b and c (b) Representative flow cytometry profiles of dissociated 5dpf p21:GFP zebrafish and wild type siblings treated with either 0Gy or 12Gy irradiation and quantitation of the proportion of live p21:GFP-, p21:GFP^Dim^, and p21:GFP^Bright^ cells in dissociated 5dpf *p21*:GFP zebrafish larvae. Dissociated cells from 50 fish were pooled for each repeat (n=3)(c) Representative flow cytometry profiles of dissociated 12dpf *p21*:GFP zebrafish larvae and wild type siblings treated with either 0Gy or 12Gy irradiation. Quantification of the proportion of live p21:GFP-, p21:GFP^Dim^, and p21:GFP^Bright^ cells in dissociated 12dpf*p21*:GFP zebrafish larvae. Dissociated cells from 25 fish were pooled for each experiment (n=3). Data were examined by 2way ANOVA with Šidak’s multiple comparison test. (d) The proportion of p21:GFP^Dim^, and p21:GFP^Bright^ at 18, 28 and at least 36 months (mo) old in adult *p21:GFP* zebrafish brains, intestines and livers were quantified. Data were examined by one way ANOVA with Sidak’s multiple comparison test. Data are presented as mean ± SEM. *** p<0.001, ** *p* < 0.01; * *p* < 0.05.

### P21:GFP^bright^ cells show multiple markers of senescence

To verify that p21:GFP^+^ cells co-expressed other known markers of senescence, p21:GFP zebrafish larvae were exposed to 12Gy irradiation at 2dpf and p21:GFP^+^ cells were subjected to fluorescence activated cell sorting (FACS) at 5 and 12 dpf. Cells showed over 90% purity following FACS sorting (Fig 5a). Following irradiation, p21:GFP^bright^ but not p21:GFP^dim^ cells, showed a significant increase in size and granularity, which are features of senescence, compared to those not exposed to irradiation (Fig 5b-d). In addition, a significant increase in cells with >5 γH2AX^+^ foci and PCNA negative was observed in both GFP^Bright^ and GFP^Dim^ populations at 5 dpf and persisted at 12 dpf (Fig 5e-g). However, the increase was more modest in the GFP^dim^ (31.2±10.1% at 5dpf) in comparison to the GFP^bright^ population (52.2±8.1% at 5dpf)(Fig 5g). No significant increase in the number of cells IL6^+^PCNA^−^ was observed in the GFP^dim^ population when compared to cells from non-irradiated p21:GFP fish at 5dpf and 12dpf (Fig 5g). This was in contrast to the GFP^bright^ population where 22.0± 3.5% of GFP^bright^ cells were IL6^+^PCNA^−^ at 5dpf and this increased to 48.1±6.7% at 12 dpf (fig. 5g). These data suggest that the p21GFP^bright^ population enriches for a population with properties of senescent cells.

**Figure 5.**
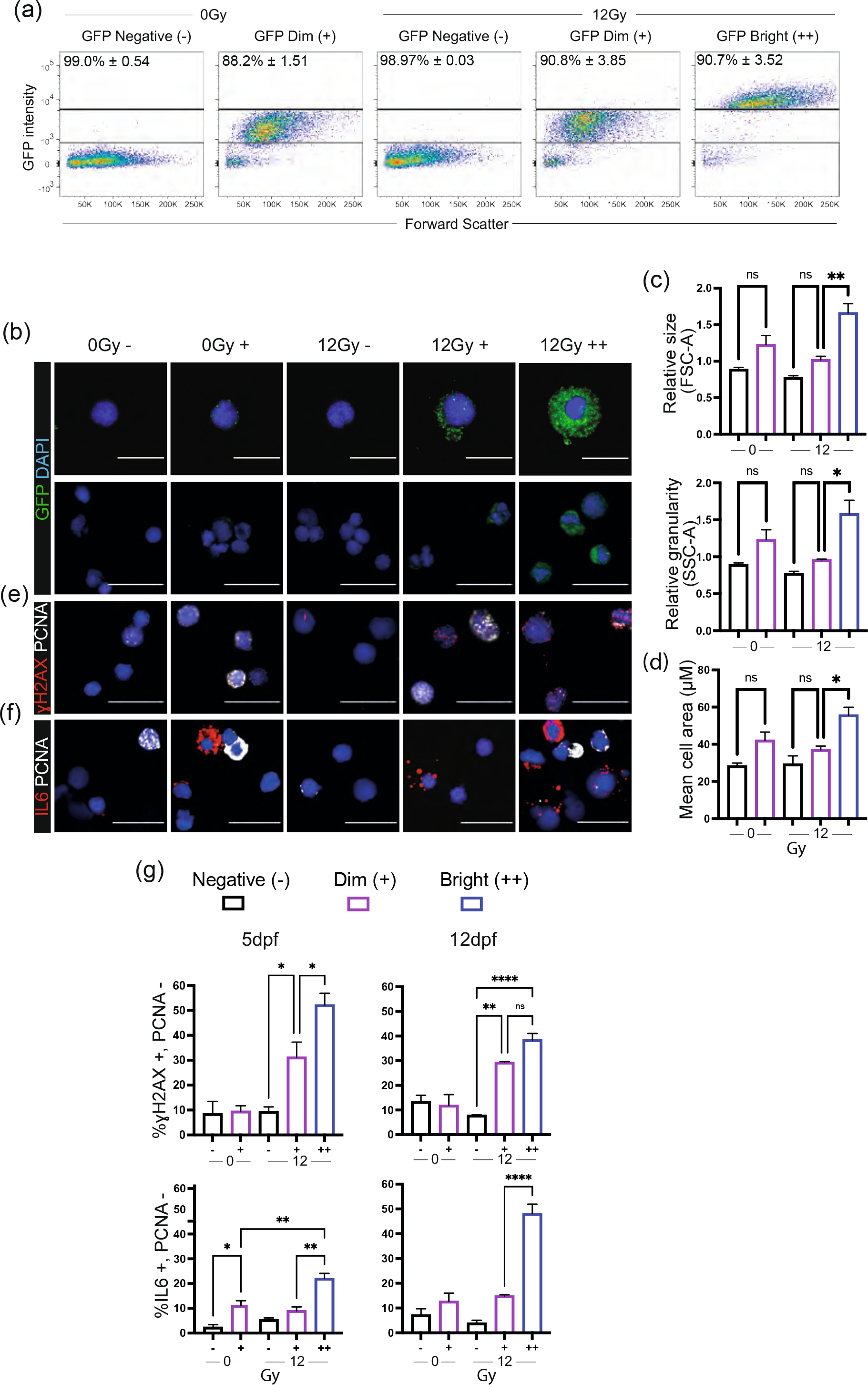
GFP^Bright^ cells are associated with other markers of senescence at 5 and 12dpf. (a) Representative flow cytometry profiles of 5dpf *p21:GFP* cells after sort according to GFP intensity. The level of purity of each population across three independent biological replicates is at the top of the graph. (b) Representative confocal fluorescent photomicrographs of immunofluorescence for GFP Scale 10μm (top), 20μm (bottom). (c) Quantification of cell size (FSC-A) and granularity (SSC-A) in GFP^Negative^, GFP^Dim^, and GFP^Bright^ populations, relative to total live cells. (d) Mean cell area quantified by measuring confocal fluorescent photomicrographs of sorted GFP^Negative^, GFP^Dim^, and GFP^Bright^ populations. (e-f) Representative confocal fluorescent photomicrographs of immunofluorescence for (e) ɣH2AX and PCNA or (f) IL6 and PCNA in 5dpf p21:GFP cells, sorted according to GFP intensity. f) Quantification of the proportion of ɣH2AX +/PCNA– cells (top) and IL6 +/PCNA– cells (bottom) in GFP^Negative^, GFP^Dim^, and GFP^Bright^ populations at 5dpf (left) and 12dpf (right). Data were examined by one-way ANOVA with Sidak’s multiple comparisons test. 300 cells quantified for each group over 3 independent experiments. Scale 20μm. Mean ± SEM represented throughout. **** *p < 0.0001*; ** *p* < 0.01; * *p* < 0.05.

### Senolytics clear p21^bright^ cells in zebrafish

To develop an accurate way to detect p21:GFP^bright^ cells by imaging for drug testing, zebrafish were irradiated at 12Gy at 2dpf and transferred in a 96-well plate in medium. At 5dpf, zebrafish were anaesthetised and imaged in a horizontal position on an Opera Phenix® High-Content Screening System. Tiled confocal photomicrographs were acquired and individual cells segregated for analysis (Fig 6a). The mean fluorescence intensity of individual cells was classified against thresholds to determine whether they were p21:GFP^−^, p21-GFP^Dim^ or p21:GFP^Bright^. The thresholds were established based on the level of fluorescence in the untreated wild type zebrafish (p21:GFP^−^) and non-irradiated p21:GFP zebrafish (p21:GFP^dim^). To verify that these thresholds detected the correct proportion of p21:GFP^bright^ in a reproducible manner, we firstly analysed p21:GFP fish at 5dpf in three experiments performed on three different days. We compared the reproducibility in detecting the zebrafish area considered for analysis, the number of fluorescent cells detected in each zebrafish and the proportion of p21:GFP^bright^/zebrafish in the whole fluorescent population. No significant difference was observed when the same plate was analysed on different days by the same operator (extended Fig 2). In addition, the number of p21GFP^bright^ cells in irradiated and non-irradiated fish was compared. As expected, we observed a significant increase in the p21:GFP^bright^ cells following irradiation (Fig 6b). Finally, the number of p21:GFP^bright^ detected by imaging were compared to those detected by flow cytometry. No significant difference was found when comparing the percentage increase in GFP^bright^ cells detected by this method and by FACS (Fig 6b), suggesting that this method is accurately measuring the increase in the number of p21:GFP^bright^ cells. To verify that known senolytics had similar effects in zebrafish to those reported in *in vitro* and *in vivo* models, p21:GFP fish were irradiated at 2dpf and transferred to a 96-well plate in medium containing either the senolytic cocktail dasatinib (D, 500nM) and quercetin (Q, 50μM) or ABT-263 (navitoclax, 5μM). DMSO was used as a vehicle control. These doses were established as the highest doses that did not cause significant acute toxicity to the fish based on key signs including pericardial oedema and abnormal spinal curvature [23]. Media containing the drugs was refreshed at 4dpf and fish were analysed at 5dpf. A decrease in the percentage of GFP^bright^/total number of fluorescent cells was observed when DQ and ABT263 were administered, although this reached statistical significance only for DQ (Fig 6c).

**Figure 6.**
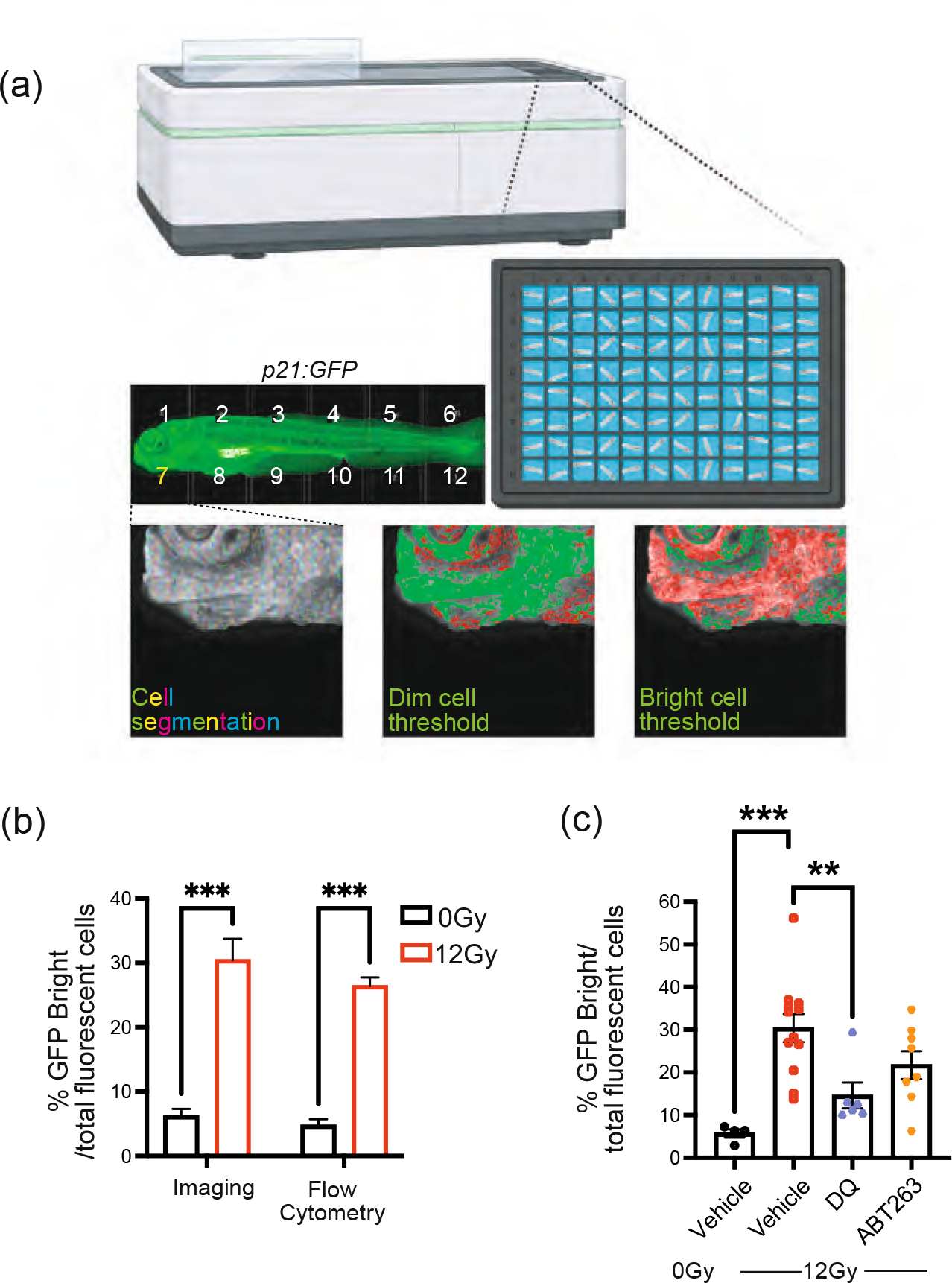
Senolytics reduces the number of p21:GFP^Bright^ cells. (a) Diagram representing automated imaging method for p21:GFP zebrafish using The Opera Phenix High-Content Screening System. Tiled confocal photomicrographs from Opera Phenix microscope of 5dpf p21:GFP zebrafish were acquired, and individual cells were segregated for analysis. The mean fluorescence intensity of individual cells was classified against a threshold set on the basis of level of fluorescence in wild type fish and non-irradiated p21:GFP fish to identify the GFP^Bright^ population; (b) Percentage of GFP^Bright^ cells calculated by Opera Phenix High-Content Imaging and Flow Cytometry analysis, as a proportion of total fluorescent cells in p21:GFP fish with or withour irradiation. (c) Quantification of the proportion of GFP^Bright^ at 5dpf in laterally-oriented p21:GFP fish following irradiation at 2dpf and treatment starting at 3dpf with vehicle, dasatinib (D) plus quercetin (Q) or ABT263 (navitoclax). Data were examined with one-way ANOVA with Tukey’s multiple comparison’s test. Data (from two independent experiments are presented as Mean ± SEM presented. *** *p* < 0.001; ** *p* < 0.01.

## Discussion

In this study we have generated a p21:GFP zebrafish model and developed a protocol for the induction of senescence over 5 days. p21 is an important marker of senescence as shown by studies in p21GFP mice [11]. As was recently found in these mice, we have identified a population of p21GFP^bright^ cells, which accumulate in both zebrafish larvae following irradiation and in the tissues with age and are enriched for markers of senescence. Approximately 50% of the GFP^bright^ PCNA- cells show more than 5 γH2AX foci/cell and express IL6 at 5dpf and at 12dpf, suggesting that they are persistent. This is comparable to what has been found in the p21GFP mice where approximately 50% of cells were found to be positive for SA-β-Gal staining [11]. The percentage of p21GFP^bright^ cells in the tissues of adult and geriatric zebrafish are similar to the level of senescence reported in the literature in other species. For example we have found about 0.8% of p21GFP^bright^ cells in geriatric fish brain and a study in human brains with various levels of Alzheimer’s disease found less than 2% of cells were senescent [16]. We have identified approximately 4% of GFP^+^ cells in zebrafish liver at 18 months of age. This was similar to the findings of Ogrodnik *et al*, 2017 in the liver of mice at a similar age [17].

We have established a rapid protocol for the induction of senescence in zebrafish larvae using irradiation. This is a well-known method for rapid induction of senescence in cells and it has been shown to accelerate signs of ageing such as frailty in mice [24]. Previous work in zebrafish showed that irradiation at 1dpf led to high levels of mortality and developmental abnormalities, similar to our findings [25, 26]. However, we have identified a treatment window using 12Gy at day 2, which does not cause significant acute toxicity and death. Expression of senescent marker p21 and SA-β-gal could be detected by 5dpf and was particularly strong in the pharyngeal arches, brain, and intestinal regions. This was similar to findings of Kishi *et al* (2008) and Silva-Álvarez *et al* (2020) [14, 27] during the identification of mutants expressing higher levels of SA-β-Gal. The mutant genes were involved in regulation of lifespan and telomere length regulation, and showed accelerated signs of ageing in adult life, suggesting the presence of *bona fide* senescence [27]. Whilst this protocol is convenient to study mechanisms of stress-induced senescence, there are other mutants available, including a *tert^−/-^* zebrafish line to study telomere induced senescence [21]. Oncogene induced senescence can be studied by injecting Ras^G12V^ cells in the zebrafish epithelia, in larvae [28], increasing the potential use of this model.

We have chosen to develop the model in zebrafish at the larval stage due to the many advantages that this model offers to test gene function or screening new compounds, and which are complementary to those of mammalian systems. As well as high fecundity and ready genetic manipulation, most organs are developed by 48-72h. By 96h the pancreas, liver and gallbladder are developed and by 120h the development of the gastrointestinal system is complete [29, 30]. Most organs perform a similar function to the human counterpart with well-conserved physiological mechanisms [30]. This means that it is possible to obtain important information on organ function, mechanisms of action and on efficacy and toxicity of compounds. When compared to *in vitro* cell testing, the model has the added value, of taking into consideration the complexities of interactions at the whole organism level. A number of tests are available to assess organ function. We have shown that larvae lose locomotor function with irradiation and there are changes in muscle fibres, which resemble those found in geriatric fish. Locomotion is not just the result of muscle function but requires an integrated response involving brain function, the nervous system and visual acuity. There are other tests available to monitor the fitness of the major organ systems, including heart, memory and cognition, liver, kidney, immune and sensorial function [30].

Their small dimensions means that each fish can easily fit in a 96-well plate, making any test relatively easy and inexpensive, requiring only small amounts of drugs. For these reasons, use of zebrafish for *in vivo* drug testing and toxicology is increasing. Whilst it is acknowledged that there are problems of poor solubility with some compounds and it is difficult to compare how toxic in-water dosing relates to mammalian plasma levels, ways to measure absorption, distribution, metabolism and excretion (ADME) in zebrafish are in development [31]. There is good agreement with findings in mammalian developmental toxicity studies reaching up to 85 [32] or 87% [33] agreement. Eight molecules are undergoing clinical testing following a combination of human genetic data and testing in zebrafish models without additional animal testing [34]. There has been an acceleration in technical development to use this model for drug testing in the last 15 years, with development of many animal models of diseases, transgenic lines, tests for cardiac, nephron and liver toxicity [34], which makes it a real promise for the future.

The transparency at the larval stage and the availability of reporter lines means that it is possible to isolate cells by flow cytometry or image them at the single cell level in the living organism, opening up opportunities for mechanistic studies in senescence. There are over 8000 transgenic lines with fluorescent reporters, which model specific diseases, label molecules, specific organelles or specific cell types, allowing their visualisation and tracking *in vivo* [35]. For example there are transgenic lines labelling most cell types of the immune system [36]. This opens opportunities to visualise and track in real time over a 24h period the interaction of senescent cells with immune cells in steady state or during regeneration. It will allow us to answer fundamental questions as to whether immune cells are responsible for the elimination of senescent cells, whether this ability is reduced with age and what is their relative contribution to the accumulation of senescent cells with age. We have demonstrated that, using this model, it is possible to identify p21:GFP^bright^ cells using imaging of individual fish in 96-well plates and at the single cell level, which reflect values observed by flow cytometry with good reproducibility. Both treatments, DQ and navitoclax induced a reduction in p21:GFP^bright^ cells at doses in the same range of what was previously published in cells i*n vitro*, although navitoclax was less effective and did not reach statistical significance. This is similar to previously published findings in cells in other species. Quercetin showed senolytic properties at 10-20μM in human adipocytes and endothelial cells and at 100 μM in mouse mesenchymal stem cells [37]. Dasatinib was given at concentrations ranging from 100-300nM in human adipocytes and endothelial cells and 500nM in mouse mesenchymal stem cells [37][33]. For Navitoclax there was a narrower range of concentration available before induction of toxicity. The biggest difference between non-senescent and senescent cells was observed at higher concentrations than the one we could use in zebrafish larvae (5μM vs 10-20μM [38]). In addition, Cai et al. (2020) compared the senolytic activity of DQ and navitoclax and senolytic effects were observed in human embryonic fibroblasts and human umbilical vein cells but not in pre-adipocytes with navitoclax [38]. This was in contrast to DQ, which was effective in all cell types although with different intensity. The reduced toxicity of DQ combined with a larger spectrum of cells affected may explain the increased effectiveness of DQ in our model. Indeed navitoclax is best when given at lower doses for a longer time to reduce its toxic effects. Its toxicity is well recognised and new compounds targeting selectively senescent cells are in development to overcome this problem [39].

In summary, we demonstrate that the p21:GFP model in zebrafish larvae offers a powerful new tool that could be used to accelerate the study of mechanisms of senescence, its relationship with disease and for drug testing purposes, allowing the selection of only the most promising mechanisms and compounds for study in mammalian models.

## Material and methods

### Husbandry and irradiation

Zebrafish were housed in accordance with the UK Home Office Licence animal care protocols in the Bateson Centre at The University of Sheffield, UK under standard conditions [40]. Procedures in zebrafish older than 5dpf were approved by the Home Office (Project License 70/8178). Animals were sacrificed by a schedule 1 method. For other procedures requiring anaesthesia 168mg/l of MS222 (Sigma, MO, USA) was used. For irradiation, zebrafish were removed from their chorions at 2dpf, placed in E3 media (5mM NaCl, 0.17mM KCl, 0.33mM CaCl2, 0.33mM MgSO4, 0.00001% Methylene Blue) and exposed to Cesium-137. After irradiation, zebrafish received fresh E3 media and returned to a 28°C incubator.

### Generation of p21: GFP transgenic line

To generate the p21:GFP reporter transgenic line we used the DKEY 192-O24 bacterial artificial chromosome (BAC), as per standard protocols [20]. This is a pIndigo BAC-536 vector encompassing the *p21* locus and containing 100kbp downstream and 590kbp upstream sequence from the start codon (ZFIN, 2019) [41]. A plasmid containing GFP and a kanamycin resistance cassette, (generated by Dr. Stone Elworthy, The University of Sheffield), amplified with Ultramer DNA oligos (IDT, IA, USA)(sequences in Table 2) was used to insert GFP directly after the *p21* locus. A tol2-transposon mediated system was incorporated according to standard protocols [20] to improve transmission efficiency. This involves modifying the GFP-BAC again through bacteriophage-mediated homologous recombination using an itol2kan plasmid (generated by Renshaw lab, The University of Sheffield) (Table 2). Finally, the modified plasmid was purified using a Nucleobond PC100 kit (Machery-Nagel, Deutschland) as per manufacturers’ instructions and quantified. Injections of the modified BAC into single-cell stage nacre-strain zebrafish embryos (<30 minutes post-fertilisation) were carried out. Injected larvae (F0s) were screen for transient GFP expression and raised to identify a stable F1 transgenic line.

### Assessment of senescence in whole fish

*In situ* hybridisation was performed according to standard protocol [42]. For *p21 in situ* hybridisation, the antisense RNA probe for *p21* was synthesised from linearised plasmid DNA provided by David Whitmore (University College London) [15]. Imaging was performed by placing larvae in 80-100% glycerol solution on a glass cover slip (Scientific Laboratory Supplies, UK). Transmitted light imaging was performed using a Nikon SMZ1500 stereomicroscope with a Prior Z-drive and a Nikon DS-Fi1 colour camera with NIS elements software (Version 4.3). For quantitation, colorimetric analysis was chosen by selection of the zebrafish head, excluding the eye and extending as far as the optic vesicle. Blind ranking was carried out, with images ordered by eye according to staining intensity whereby the strongest staining had the highest rank.

To assess SA-β-Gal activity in zebrafish larvae, the procedure was carried out as previously described [27]. Briefly, zebrafish were fixed in 4% paraformaldehyde in PBS overnight. Larvae were then washed for 1 hour in PBS at pH 7.45 and then for 1 hour in PBS at pH 6.0 at 4°C. The enzymatic reaction was then performed overnight at 37°C in 5mM potassium ferrocyanide, 5mM potassium ferricyanide, 2mM MgCl_2_, and 1 mg/ml X-gal in PBS (pH 6.0) (Cell Signalling Technology). Afterwards, zebrafish were fixed in 4% PFA for 4 hours and imaging and quantification were performed as described for p21 *in situ* hybridization analysis.

For γH2AX staining in the whole zebrafish, after fixation, fish were then transferred to glass reaction vials and incubated with acetone for 7 minutes at −20°C. Fish were then washed twice with PBS-T (PBS, 0.1% Tween-20, Sigma-Aldrich, UK) and blocked for 60 minutes in PBS-T with 2% blocking reagent (Roche, UK), 5% foetal calf serum (Sigma-Aldrich, UK) and 1% DMSO (Sigma-Aldrich, UK). Zebrafish were then incubated with histone H2A.XS139ph (phospho Ser139) antibody (Genetex, CA, USA, 1:1000 dilution in blocking solution), washed and incubated with Alexa Fluor 568 goat anti-rabbit (Invitrogen, UK, 1:500 dilution in blocking solution) for two hours at room temperature in the dark. Following further washing on PBS-T DAPI (Sigma-Aldrich, UK, 1:2000 dilution in PBS) was added for 15 minutes at room temperature and the fish was mounted with Vectashield (Vector laboratories, UK). The UltraVIEWVoX spinning disc confocal laser imaging system (Perkin Elmer) was used to image the tails or heads of these fish, using 1μM z-stacks with the Prior 200μm z-piezo. Images were analysed on VolocityTM software (version 6.3) to determine the number of cells positive for pγH2AX. The number of nuclear bright red+ cells were enumerated in 4 regions of interest. At least 300 cells were quantified for each test.

### qPCR

Zebrafish embryos were homogenised by addition of Trizol (Sigma, USA) and passed through a QIAshredder column (Qiagen, Netherlands) to further homogenise the tissue. The RNA was then purified through chloroform (Sigma, USA), and isopropanol (Sigma, USA) precipitation and First-Strand cDNA Synthesis was performed using SuperScriptTM II Reverse Transcription kit (Invitrogen, UK) as per manufacturer’s instructions. qPCR was performed using MESA GREEN qPCR MasterMix Plus for SYBR® (Eurogentec, UK) and an ABI 7900HT Sequence Detection System (Applied Biosystems, CA, USA). Quantification was carried out by fold expression change following irradiation (2^-ΔΔCt, relative to βactin and unirradiated control). Primers sequences are listed in Table 1.

**Table 1.**
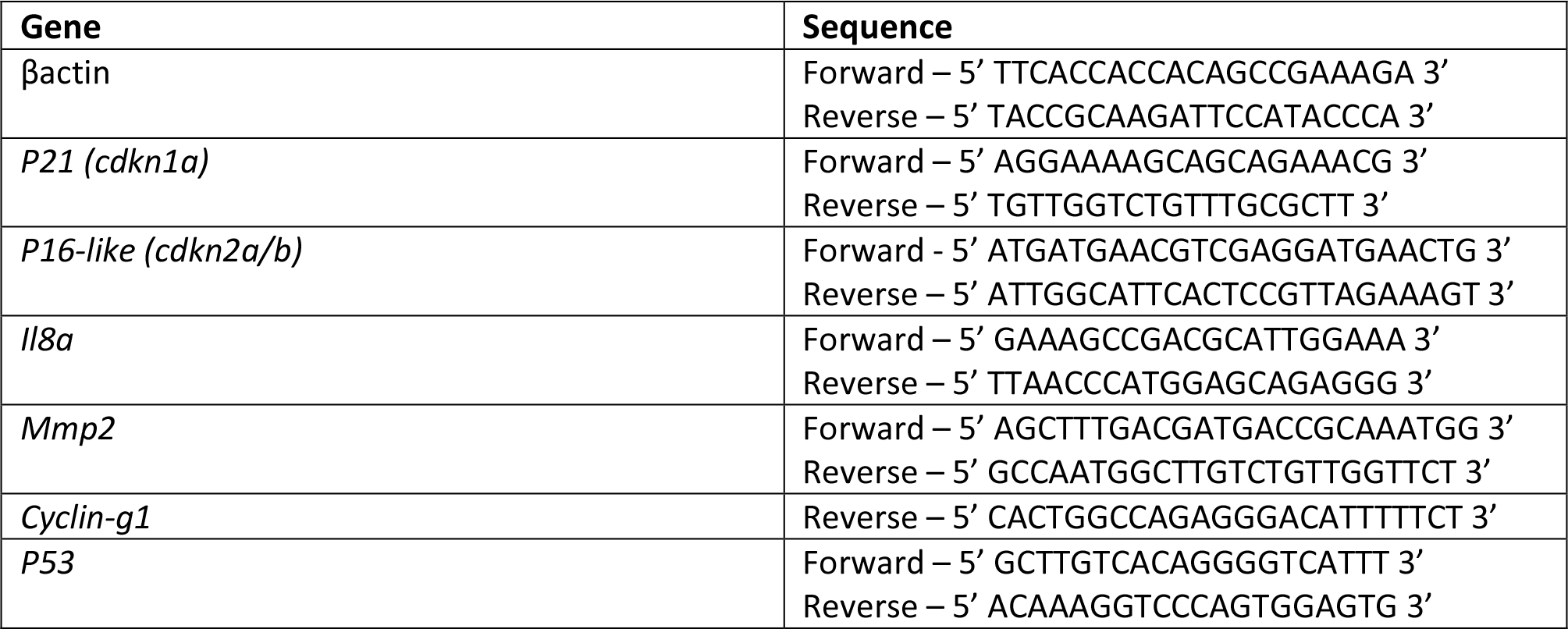
Table of primer sequences for qPCR.

**Table 2.**
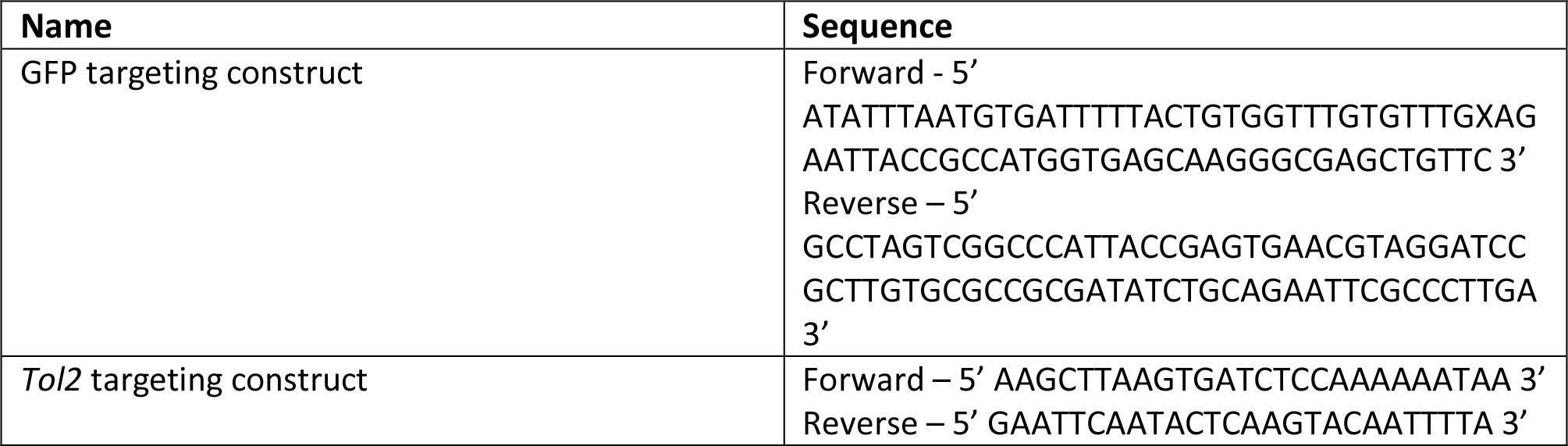
Primer sequences used to modify DKEY-192O24 BAC.

### Histology of muscle

Adult zebrafish were fixed in 10% buffered formalin for 72 hours at 4°C before decalcification in 0.5M EDTA for 72 hours at 4°C. Zebrafish larvae were fixed in 10% buffered formalin for 24 hours without decalcification. Fixed zebrafish were then paraffin-embedded and sectioned longitudinally at 3μm thickness. Zebrafish sections were stained with Hematoxylin-Eosin for histopathological analysis. To analyse the H&E-stained zebrafish, slides were digitised by a Pannoramic 250 Flash II slide scanner (3D Histech, Hungary) at 20× magnification. Zebrafish muscle width was then quantified across the larval tail, or ventral muscle for adult zebrafish.

### Analysis of locomotion

Locomotor activity was recorded using the Zebrabox (ViewPoint Life Sciences, Lyon, France) tracking system and Zebralab software (ViewPoint Life Sciences, Lyon, France). Zebrafish were placed in 24-well plate in E3 medium and movement was recorded for 30 min. Light and dark cycles of 5 minutes at 100% light and 5 minutes at 0% light were used to stimulate movement of the fish. Total movement across the 30-minute period was assessed with Zebralab.

### Fluorescent activated cell sorting

Zebrafish larvae were digested with Liberase TL (Roche, UK) at 40μg/ml and incubated at 37°C for 35 minutes. This was followed by addition of 10% Fetal Bovine Serum (FBS) (Sigma-Aldrich, UK) and passed through a 40μm mesh. The suspension was then centrifuged at 1,500rpm for 5 minutes, before cells were re-suspended in Leibovitz’s L15 media (ThermoFisher Scientific Inc., USA), containing 20% FBS and 5mM EDTA. Cells were incubated with 7-Aminoactinomycin D (7AAD, ThermoFisher Scientific Inc., USA) for 5 minutes before being examined by a LSRII (BD) flow cytometer (BD Biosciences, San Jose, CA, USA). For the assessment of adult zebrafish organs, they were dissected and manually dissociated with a sterile scalpel before being digested as described above for the zebrafish larvae.

### Immunofluorescent staining of single cells

Following fluorescent activated cell sorting, p21:GFP cells were washed in PBS and fixed with 4% paraformaldehyde for 20 minutes on ice. Cells were then cytospun at 500rpm for 5 minutes with medium acceleration onto SuperFrost Ultra Plus™ adhesion slides (ThermoFisher Scientific Inc., USA) and dried overnight at room temperature (22°C). Cells were then permeabilised with 0.5% Triton X-100 (Sigma-Aldrich) for 10 minutes at room temperature and blocked for 1-2 hours at room temperature with 3% Bovine Serum Albumin, 5% Goat Serum, 0.3% Tween-20 in PBS. Slides were then incubated with primary antibodies overnight at 4°C. A Combination of antibodies against Proliferating Cell Nuclear Antigen (PCNA, Santa Cruz, CA, USA, 1:200 dilution) and histone H2A.XS139ph (phospho Ser139) antibody (Genetex, CA, USA, 1:300 dilution) or Proliferating Cell Nuclear Antigen (PCNA, Santa Cruz, CA, USA, 1:200 dilution) and IL6 (Abcam, MA, USA, 1:500 dilution) were used. This was followed by overnight incubation with secondary antibodies at 4°C in blocking solution. Combinations of antibodies against Alexa Fluor 488 goat anti-chicken (Abcam, MA, USA, 1 in 500 dilution), Alexa Fluor 568 goat anti-rabbit (Invitrogen, UK, 1:500 dilution) and Alexa Fluor 647 goat anti-mouse (Invitrogen, UK, 1:500 dilution) were used. Slides were stained with DAPI (Sigma-Aldrich, UK, 1:2000 dilution in PBS) and then mounted in Vectashield (Vector laboratories, UK). Slides were imaged on a Deltavision microscope using an UplanSApo 40x oil objective (NA 1.3) and Photometrics CoolsnapHQ CCD camera. Z stacks were imaged at 1μm and deconvolution software was used for the pγH2AX staining to make the foci more visible. Excitation by a 100W Hg lamp was used. Quantification was carried out with maximum intensity projections of 15 z-stacks (15μm thickness). Cells with 5 or more γH2AX foci in a single nucleus were deemed positive (stained with DAPI). Cells positive for IL6 had clear red fluorescence in the regions around the nucleus, whilst cells positive for PCNA had clear nuclear staining. At least 300 cells were quantified per group.

### Opera Phenix imaging

Zebrafish were dechorionated and irradiated as previously described at 2dpf. At 3dpf, zebrafish were placed in individual wells of a 96-well μclear® cell culture imaging plates, containing either the senolytic cocktail Dasatinib (D, 500nM) and Quercetin (Q, 50μM) or ABT- 263 (Navitoclax, 5μM), dissolved in 0.5% DMSO which was used as vehicle control. At 4dpf, the drug media was refreshed. At 5dpf, zebrafish were anaesthesised and imaged on an Opera Phenix® High-Content Screening System (Perkin Elmer). The 96-wells were first imaged at 5x magnification to identify zebrafish via their GFP intensity, before a 20x high magnification z-stack was taken of the whole zebrafish larvae. Images of 5dpf zebrafish were quantified on Harmony® High-Content Imaging and Analysis Software. To first identify our region of interest, the whole zebrafish, the ‘Common Threshold’ method was used on the Alexa 488 channel to detect GFP intensity. A threshold of 0.25 was used. This was used to determine the mean fluorescence intensity of p21:GFP zebrafish. To identify and segregate cells on the detected zebrafish, method ‘M’ was utilised with a diameter of 20uM, splitting sensitivity at 0.48, and common threshold at 0.22. The intensity properties of individual cells are analysed to threshold GFP^Dim^ and GFP^Bright^ cells. GFP^Bright^ cells have a mean Alex 488 intensity of at least 1250, while dim cells have an intensity between 400 and 1250. This was established using untreated wildtype zebrafish, 0Gy (mock) and 12Gy irradiated zebrafish, before results were compared to flow cytometry data to ensure the proportion of GFP^Dim^ and GFP^Bright^ cells were comparable with both techniques. Next, the number of GFP^Dim^ and GFP^Bright^ cells in the 5dpf zebrafish was automatically counted and determined to establish whether the senolytics were able to selectively remove fluorescent cells from p21: GFP zebrafish.

### Statistical analysis

Data were analysed by Prism software (version 8.1). Data were analysed by t-test or one way ANOVA. Analysis and post-hoc tests carried out for individual data sets are specified in their respective figure legends with significance *P < 0.05, **P < 0.01, ***P < 0.001, ****P < 0.0001.

## Acknowledgements

Samir Morsli was funded by the Biotechnology and Biological Sciences Research councils PhD studentship (DTP2 BBSRC Grant Ref: BB/M011151/1). This research (Heather Mortiboys) was supported / funded by the NIHR Sheffield Biomedical Research Centre (BRC) / NIHR Sheffield Clinical Research Facility (CRF). The views expressed are those of the author(s) and not necessarily those of the NHS, the NIHR or the Department of Health and Social Care (DHSC). Figures 3A & 6A were created with BioRender.com. We are grateful to Stone Elworthy for providing the GFP plasmid and David Whitmore (University College London) for providing the p21 probe.

