## Extended Fig 1 for "A p21-GFP zebrafish model of senescence for rapid testing of senolytics *in vivo*"

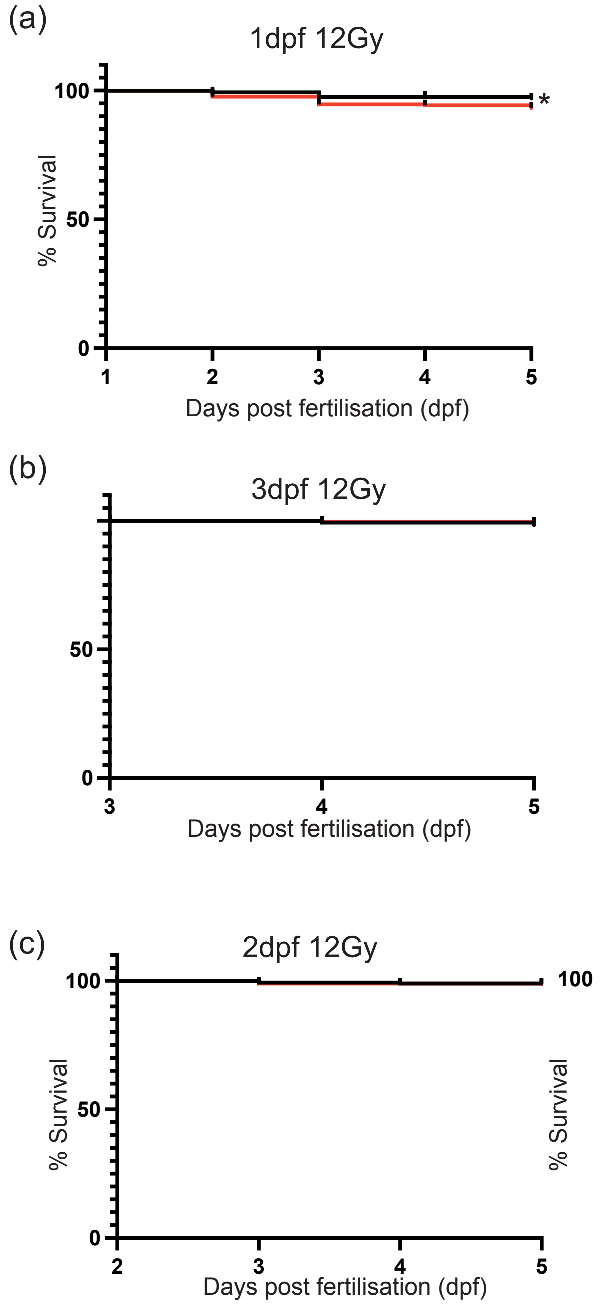


**Extended Figure 1. The dose of 12Gy Cs137 -γ-Irradiation administered at 2dpf is optimal to limit toxicity in zebrafish larvae before 5dpf.** 12Gy Cs137 -γ-Irradiation was given at 1dpf (b), 2dpf (c) or 3dpf (d) and survival examined at 5dpf by Kaplan-Meier analysis. Significant differences displayed as * $p<0.05$. Black line non irradiated fish and red line irradiated fish
