## Extended Fig 2 for "A p21-GFP zebrafish model of senescence for rapid testing of senolytics *in vivo*"

Extended Figure 2

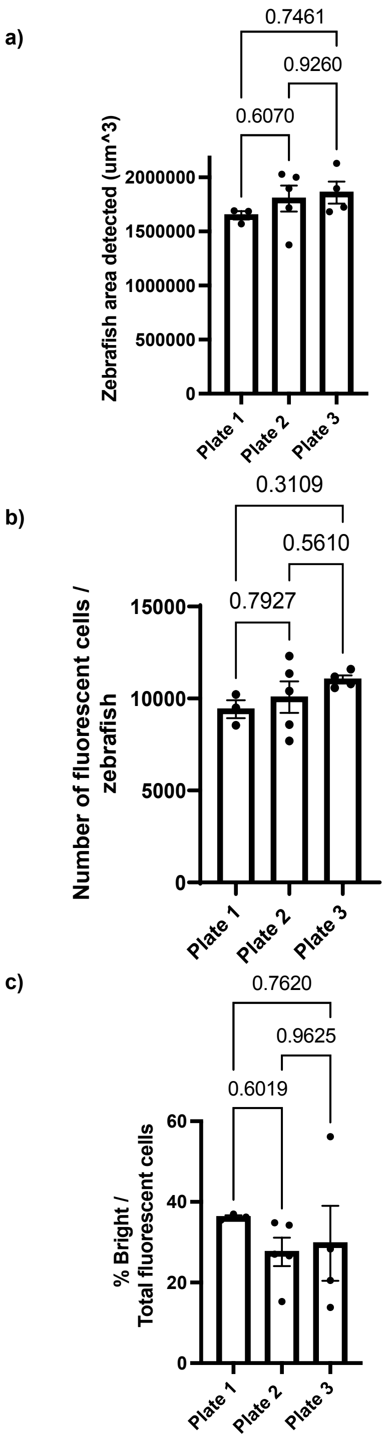


**Extended Figure 2 Opera Phenix High-Content Imaging validation of p21:GFP zebrafish.** Laterally oriented, 12Gy irradiated, 5dpf zebrafish were analysed on 3 independent days to assess assay variability. There were no detectable changes in the (A) fluorescent zebrafish area detected, (B) the number of fluorescent cells detected per zebrafish, (C) the proportion of p21:GFP$^{Bright}$ cells per zebrafish. Data were examined with one-way ANOVA with Tukey's multiple comparison's test. Mean ± SEM presented.
